# Personalized disordered circuit modeling reveals diverse neurobiological routes to transdiagnostic symptoms

**DOI:** 10.64898/2026.08.10.744027

**Authors:** Alexander J. Simon, Santino Iannone, Anja Samardzija, Sarah A. Cutts, Flor Parra, Ashlea Segal, Sydney E. Smith, Kira Y. Tang, Fuyuze Tokoglu, Jagriti Arora, Maolin Qiu, Rachel Katz, Scott Woods, Vinod Srihari, Gerard Sanacora, Xilin Shen, R. Todd Constable

## Abstract

Precision psychiatry research has strived to characterize the neural circuitry underlying transdiagnostic symptoms with the goal of devising biologically informed diagnostic and treatment strategies. However, few clinically actionable findings have emerged from these initiatives. One potential complication in reaching this goal is that similar symptoms may arise from distinct patterns of neurobiological dysfunction. Therefore, models that can provide insight into where an individual’s disordered circuits are located are important. Here, we developed a data-driven approach to identify individualized patterns of disordered connectivity using a large (*n*=317), transdiagnostic population with extensive fMRI data. Connectome-based predictive modeling identified functional connectivity networks associated with a broad array of psychiatric symptoms. Symptom networks were aggregated, and connections contributing most strongly to each individual’s symptoms were identified to construct personalized “disordered circuit models”. Within these models, many features were associated with specific diagnoses and patterns of comorbidity, despite the symptoms exhibiting no diagnostic specificity. This suggests that similar symptoms can emerge through diverse neurobiological mechanisms depending on an individual’s diagnostic profile. By characterizing patient-specific neural dysfunction, disordered circuit modeling offers a promising framework for advancing precision psychiatry by accounting for diverse mechanisms from which symptoms may arise.

---

Although categorical diagnostic frameworks, such as the DSM-5 and ICD-10, are the standard for diagnosing and determining the best course of treatment in psychiatry, patient outcomes to treatment are highly variable^1–5^. Because of this, efforts have been made to identify biomarkers that can predict treatment responses and clinical trajectories^1,4,6–8^. However, substantial symptom heterogeneity within diagnostic categories has stymied these efforts^9,10^. In recognition of this challenge, and because many symptoms are shared between diagnostic categories (e.g., sleep disturbances are common to major depression, generalized anxiety disorders, and posttraumatic stress disorder)^11–14^, the NIMH launched the Research Domain Criteria (RDoC) initiative to support investigations of the biological underpinnings of symptoms at multiple scales, including neural circuits^15,16^. Although this framework has been widely adopted within the psychiatric neuroimaging field^17–23^, the long-term goal of using findings that have emerged from this initiative to guide clinical practice has yet to be realized^8,24,25^. This may be in part because symptoms have complex interactions with each other^26^ and have heterogenous causes from person to person^2,27,28^. Furthermore, tools for precisely assessing individual specific dysfunction, such as how someone’s disordered circuits underlying their multivariate symptom profiles are distributed throughout the entire brain, are underdeveloped.

In line with the RDoC framework, network mapping techniques, such as connectome-based predictive modeling (CPM)^29,30^, have been extensively used to link functional MRI (fMRI) connectivity data to a wide range of clinically relevant phenotypes^31–40^. The resulting networks reflect where disordered circuitry related to symptomatology tend to be distributed throughout the brain. However, an individual’s disordered circuitry may not cleanly overlap with a predictive network of a particular symptom. Instead, a symptom may arise through an individual’s disordered circuitry partially overlapping with a broader network related to that symptom. Using this logic, comorbidity may then reflect whether an individual’s disordered circuitry impinges upon regions that are shared between multiple symptom networks. To this end, several studies have highlighted that shared and unique features between networks related to multiple psychiatric symptoms and mental health measures may help explain some aspects of symptom comorbidity and heterogeneity^36,40–45^. However, this information has not yet been aggregated into a unifying model that identifies where an individual’s disordered neural circuits are distributed. Such a model could greatly inform precision psychiatry practices.

Network-based models of psychiatric symptoms have garnered excitement for their potential utility as biomarkers for psychopathology^25,35,46^. However, network biomarkers that are robust enough to be implemented effectively in clinical settings have not yet been established^25^. One possible explanation why is all connections (or “edges”) within a predictive network are typically aggregated (or weighted at the group level) when examining individual differences in network connectivity. If there are multiple neurobiological mechanisms from which a symptom can arise^2,27^, then this approach will obscure the pathological features in an individual. Because predictive models are generated at the group level, it is challenging to identify whether there are meaningful variations in the edges linked to a specific symptom between individuals. There is evidence suggesting that meaningful, systematic variation exists between individuals in the edges driving network-based models of various clinically-relevant neurocognitive measures^47^. This finding implies that some of the circuit heterogeneity observed within clinical populations^48^ is not only driven by symptom heterogeneity, but also from there being multiple neurobiological pathways giving rise to similar symptoms. Failing to account for this may be preventing the field of brain-behavior modeling from yielding translationally actionable models. Understanding the relationship between an individual’s specific disordered circuits and a spectrum of networks associated with symptoms could thus provide rich, and unexplored, information. Such an endeavor would move the field away from a nomothetic approach to identifying biomarkers and toward the idiographic principle that understanding individual differences is essential for advancing precision psychiatry^49^.

In this study, we sought to characterize individual differences in the neurobiological mechanisms underlying transdiagnostic symptomatology as a function of patient’s diagnoses. To do so, we developed an approach for identifying specifically where an individual’s disordered circuits are located. Using a large (*n*=317) transdiagnostic dataset of adults with extensive fMRI scanning (>50 min from 6 tasks and 2 resting runs)^20^, we used CPM to identify network correlates of a multitude of items indexing a range of psychiatric symptoms. We then constructed ‘disordered circuit models’ that jointly accounted for an individual’s multivariate symptom profiles, the complex shared and unique network space, and edge-level heterogeneity within each symptom network. Although all symptoms spanned multiple diagnostic categories, disordered circuit modeling revealed features differentially associated with diagnostic specificity and comorbidity, revealing how distinct neurobiological mechanisms can converge on similar symptom presentations depending on an individual’s diagnostic profile. Thus, we introduce an approach for identifying individualized patterns of pathological functional connectivity that holds the potential to substantially inform precision psychiatry frameworks.

## Results

### Network architectures of diverse symptoms

Symptoms that reveal information about disordered circuitry must have clearly identifiable brain network correlates. To this end, 1000 iterations of 10-fold CPM was utilized to identify functional connectivity networks related to individual symptoms indexed by the Brief Symptom Inventory (BSI)^50^. Overall, CPM identified brain-wide networks significantly predictive of 24 different symptoms (*pFDR* < 0.05). Of these items, 3 were related to depression, 3 anxiety, 2 hostility, 3 interpersonal sensitivity, 3 paranoid ideation, 5 somatization, 1 phobic anxiety, 1 psychoticism, 1 fear, 1 sleep, and 1 guilt. The prediction strength of these symptoms ranged from *ρ* = 0.131 to 0.193 (**Fig 1a**; mean *ρ* = 0.162 +/- 0.018 stdev). Full details about the symptoms that were and were not successfully modeled by CPM can be found in **Supplemental Fig 1**. Symptom prevalence and co-occurrence may be reflected by how similarly symptom networks are related to each other. Therefore, we examined the Dice similarities between all pairs of the 24 CPM-derived symptom networks (**Fig 1b**). Correlations between these symptom scores (**Supplemental Fig 2)** were highly correlated with the network similarities (r = 0.907, p < 0.001), affirming that, because symptom networks were derived from symptoms, the network similarity largely reflects the correlational structure of the symptoms. Symptom networks showed a wide range in their similarities to each other (min Dice = 0.058; Max Dice = 0.573). The networks linked to all 3 depression symptoms were highly similar (mean Dice = 0.507). Additionally, psychoticism (mean Dice = 0.454), guilt (mean Dice = 0.444), the 3 anxiety symptoms (mean Dice = 0.343), trouble falling asleep (mean Dice = 0.277), fear (mean Dice = 0.313), and all 3 interpersonal sensitivity networks (mean Dice = 0.305) displayed notably high similarities (Dice > 0.25) to the 3 depression symptom networks. While these results highlight that these symptom networks tend to show substantial overlap, the Dice similarities ranging from 0.25 - 0.5 indicate that each network also contains unique features. This is further highlighted by **Fig 1c** showing that most edges only belong to one symptom network (7890/18630, or 42.35%), while considerably fewer are shared between multiple networks (2721 or 14.61% were shared between more than 5, and 721 or 3.87% were shared between more than 10). Consistent with previous reports, the symptom networks were distributed widely across the brain^29,36,40^ (**Fig 1d**). Overall, these results highlight that the symptom networks contain shared and unique features that span the entire brain.

**Figure 1.**
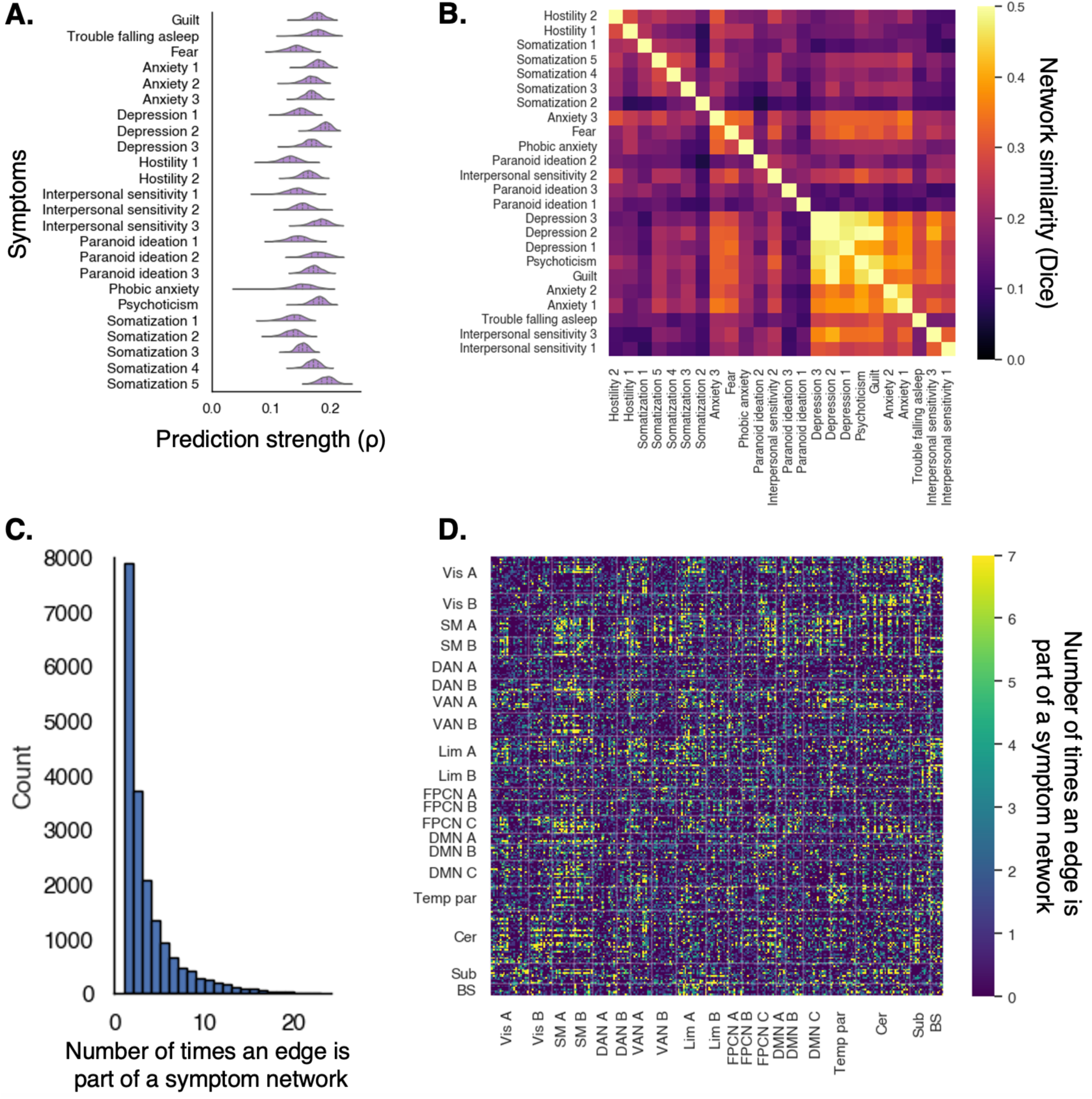
Shared and unique features of diverse symptoms. A) Distributions of prediction strength of each BSI item across 1000 CPM permutations. B) Pairwise Dice similarities of the derived symptom networks. Symptom networks were sorted by similarity based on an agglomerative hierarchical clustering algorithm. C) Histogram displaying the number of symptom networks that edges belonged to. D) Network localization of edges belonging to symptom networks were distributed throughout the brain. The binarized masks for each of the 24 symptom networks were summed and displayed to highlight where the most common edges implicated in symptomatology were.

### Symptom networks are heterogenous

We then sought to characterize the extent to which individuals varied in their edge-level dysfunction within CPM-derived symptom networks. In the CPM pipeline, the connectivity of all edges associated with a symptom in the training data are summed within an individual, and the predictive models are constructed using this sum^30^. However, if individuals differ in their neurobiological dysfunctions related to a symptom, which would be reflected by differences in the edges within symptom networks that are disordered, then aggregating the edges in a predictive model would obscure this effect. Therefore, we examined interindividual edge dysfunction heterogeneity in each of the symptom networks defined at the group level (**Fig 2a**). To do so, connectivity in every edge was residualized with respect to symptom severity to compare edge dysfunction across all participants regardless of their symptom levels. The sign of the residual-scaled edge connectivity (in z-scores) was flipped for edges that negatively predicted symptoms to ensure that higher connectivity was always more disordered. Therefore, the edges with the largest scaled connectivity values contributed most to an individual’s symptomatology.

**Fig 2.**
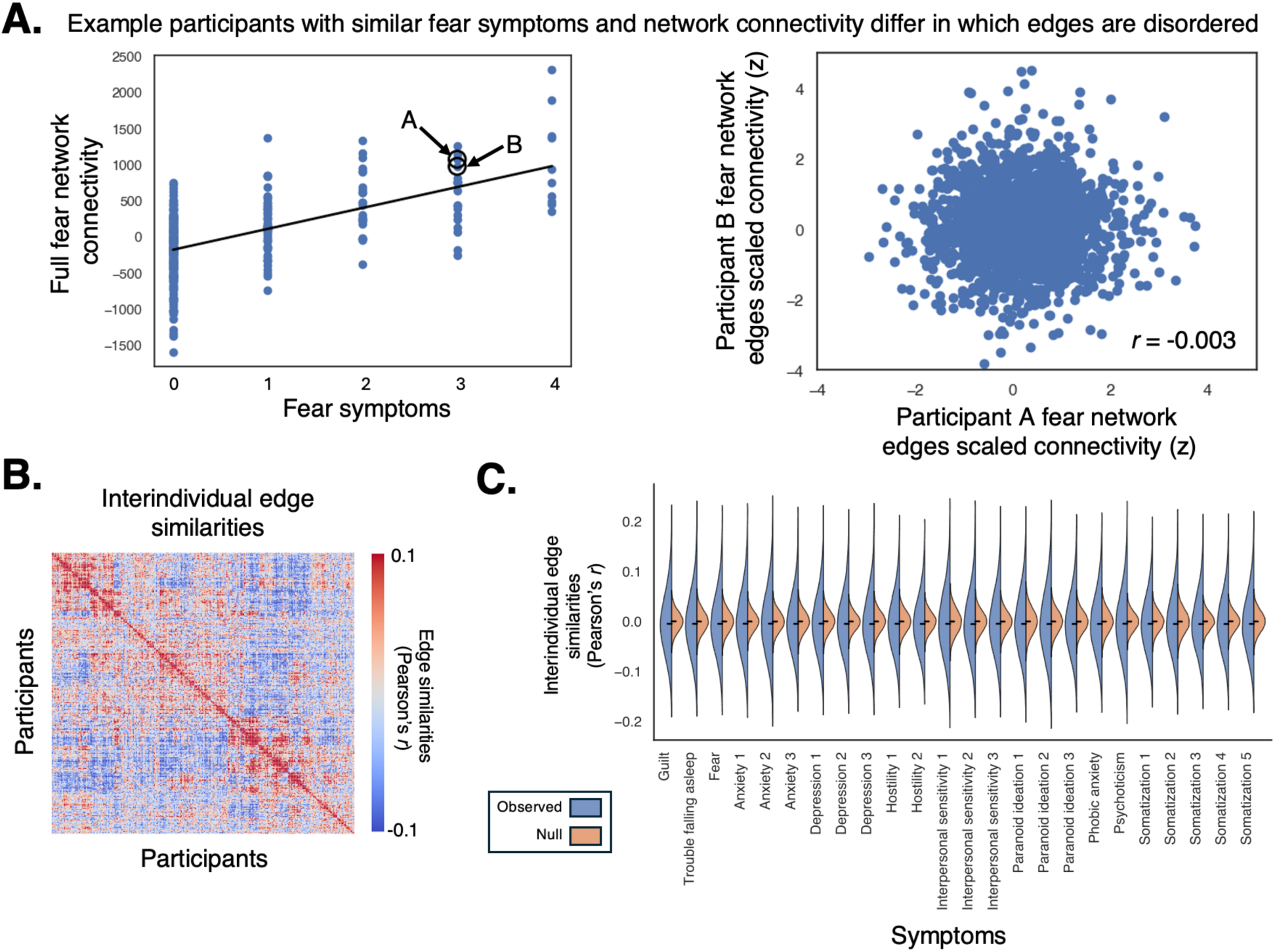
Heterogeneity in features within symptom networks. A) An example highlighting that individuals with highly similar fear symptoms can appear nearly identical when examining the connectivity of the fear symptom network as a whole, but differ dramatically in the specific edges within the symptom network that are disordered. B) A participant x participant correlation matrix showing how similar individuals were in how their scaled edge connectivities were distributed throughout an example symptom network (fear). For visualization, participants were ordered based on similarity as determined by an agglomerative hierarchical clustering algorithm. Interindividual edge-level similarity matrices for the rest of the symptom networks can be found in **Supplemental Fig 3**. C) Distributions of the participant by participant pairwise similarity (Pearson’s *r*) in scaled edge connectivity for each symptom network compared to the distribution of randomly shuffled interindividual edge similarities with the median kurtosis value.

We then investigated how edge connectivity was differentially dispersed throughout symptom networks between individuals to better understand how symptom networks were heterogeneously affected. This was achieved by correlating the scaled edge connectivities between every pair of participants for each symptom network (interindividual edge correlations in the ‘fear’ network are displayed in **Fig 2b**). Overall, interindividual similarities in edge connectivity dispersion were low (across all symptom networks: median *r* = -0.005, +/- 0.065; median and standard deviation values for each symptom network are reported in **Supplemental Table 2**). However, permutation testing revealed that there was systematic structure beyond random noise governing where disordered edges were distributed. Specifically, comparing the kurtosis of the observed interindividual edge similarity distributions to kurtosis values derived from 1000 randomly generated null distributions revealed that individuals were similar at levels significantly greater than chance (*pFDR* < 0.001) in how edge connectivity was distributed throughout each symptom network (**Fig 2c**). The kurtosis values of each symptom network can be found in **Supplemental Table 2.** These results highlight that disordered circuitry can impinge on different parts of a symptom network, leading to the same observable symptom.

### Constructing disordered circuit models

We then combined information about multivariate symptomatology with the complexities of the shared and unique symptom network space and edge-level network heterogeneity to develop a novel approach for identifying where an individual’s disordered circuits were distributed. This was achieved by first weighing each edge within each symptom network by how disordered it was for every individual (according to the residual-scaling). This step effectively enabled us to move beyond examining only the network masks and retain information about individual-specific edge heterogeneity within symptom networks. Then, each individual’s edge-weighted network masks were weighted once more by their normalized symptom severity scores. Doing so allowed us to also account for more general relationships between entire symptom networks and symptoms. Given that there is a high amount of comorbidity between symptoms and that patterns of comorbidity are variable from person to person^11,12^, values from the symptom- and residual-weighted edges were summed across all networks and normalized across all participants (z-scored). This step highlighted both the edges that were consistently implicated in poor symptomatology. This framework is presented in **Fig 3a**. Because the symptom networks partially overlapped and partially contained unique edges, after modeling 24 symptoms, we were able to identify ‘disordered circuit values’ for 18,630 out of the 35,778 total edges in the functional connectome (52.07%). The resulting disordered circuit vectors were specific to an individual’s multivariate symptom profile and their unique patterns of within- symptom network edge heterogeneity. Split-half resampling revealed that the disordered circuit vectors were highly stable, as each individual’s median correlation between the original vectors and the vectors derived from 1000 iterations of resampling ranged from 0.981 – 0.996 (median = 0.993; IQR = 0.992 – 0.994) (**Supplemental Fig 4**).

**Fig 3.**
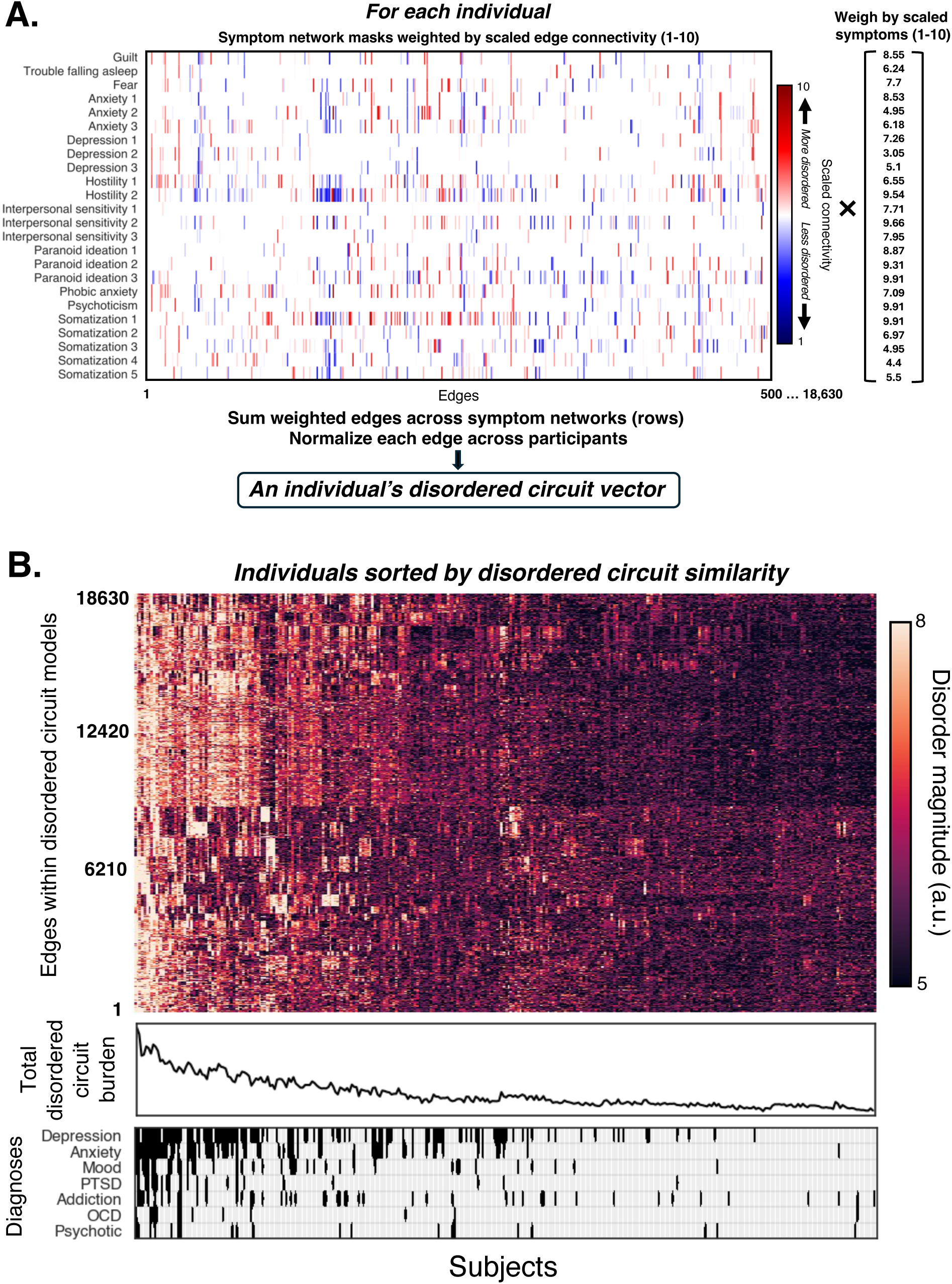
Disordered circuit models. A) The framework depicting how disordered circuit models were generated. Each row in the matrix represents a vectorized symptom network mask. The columns represent edges. For each individual, the edges within each group derived symptom network mask were weighted by the edge’s scaled dysfunction (scaled from 1-10). Then, each symptom network mask was weighted again by that individual’s symptom severity score. The doubly weighted symptom networks were then summed to highlight the edges that were consistently implicated in poor symptomatology. This was repeated for each participant, and then each edge was z-scored across participants to facilitate comparisons between edges. This yielded a disordered circuit vector for each individual. B) Individuals were sorted based on disordered circuit vector similarity using an agglomerative hierarchical clustering algorithm. Edges within disordered circuit vectors were sorted using the same approach. Participants on the left tended to have the highest burdens of disordered circuits, while participants towards the right had the lowest. There was a general correspondence between total disordered circuit burden and diagnostic load.

Once disordered circuit vectors were computed for each participant, individuals were sorted based on similarity by applying a hierarchical clustering algorithm on the disordered circuits (**Fig 3b**). This revealed a strong relationship between total disordered circuit burden (the sum of all disordered circuits for an individual) and number of diagnoses (*ρ* = 0.608, *p* < 0.001). This finding validates that disordered circuit models constructed from transdiagnostic symptoms and CPM-derived symptom networks contained information generally related to DSM/ICD diagnostic categories.

### Mapping disordered circuits to diagnostic specificity and comorbidity

We then sought to examine whether disordered circuit models contained information about the neurobiology of specific diagnoses and comorbidity patterns. To this end, multiple regression models were constructed to identify the extent to which each edge within the disordered circuit models were related to various diagnostic categories assessed by the Mini-International Neuropsychiatric Interview (MINI)^51^. After performing an FDR correction for multiple comparisons, this revealed that 7951 edges were significantly associated (*p* < 0.05) with a depression diagnosis, 14327 with an anxiety diagnosis, 8333 with a mood disorder, 6609 with PTSD, and 2962 with OCD (**Fig 4a**). None of the edges in the disordered circuit models were significantly associated with addiction or psychotic disorders. Most edges were associated with multiple diagnostic categories (mode = 2), highlighting how disordered circuits may be commonly implicated in co-morbid pathology (**Fig 4b**). We also identified edges that were specifically associated with individual diagnostic categories and with numerous combinations of diagnoses (**Fig 4c**). Notably, there were 107 edges uniquely linked to a depression diagnosis, 1796 unique to anxiety disorders, 865 unique to a mood disorder, 1296 unique to PTSD, 234 unique to OCD, and 227 that were associated with all 5 diagnoses.

**Figure 4.**
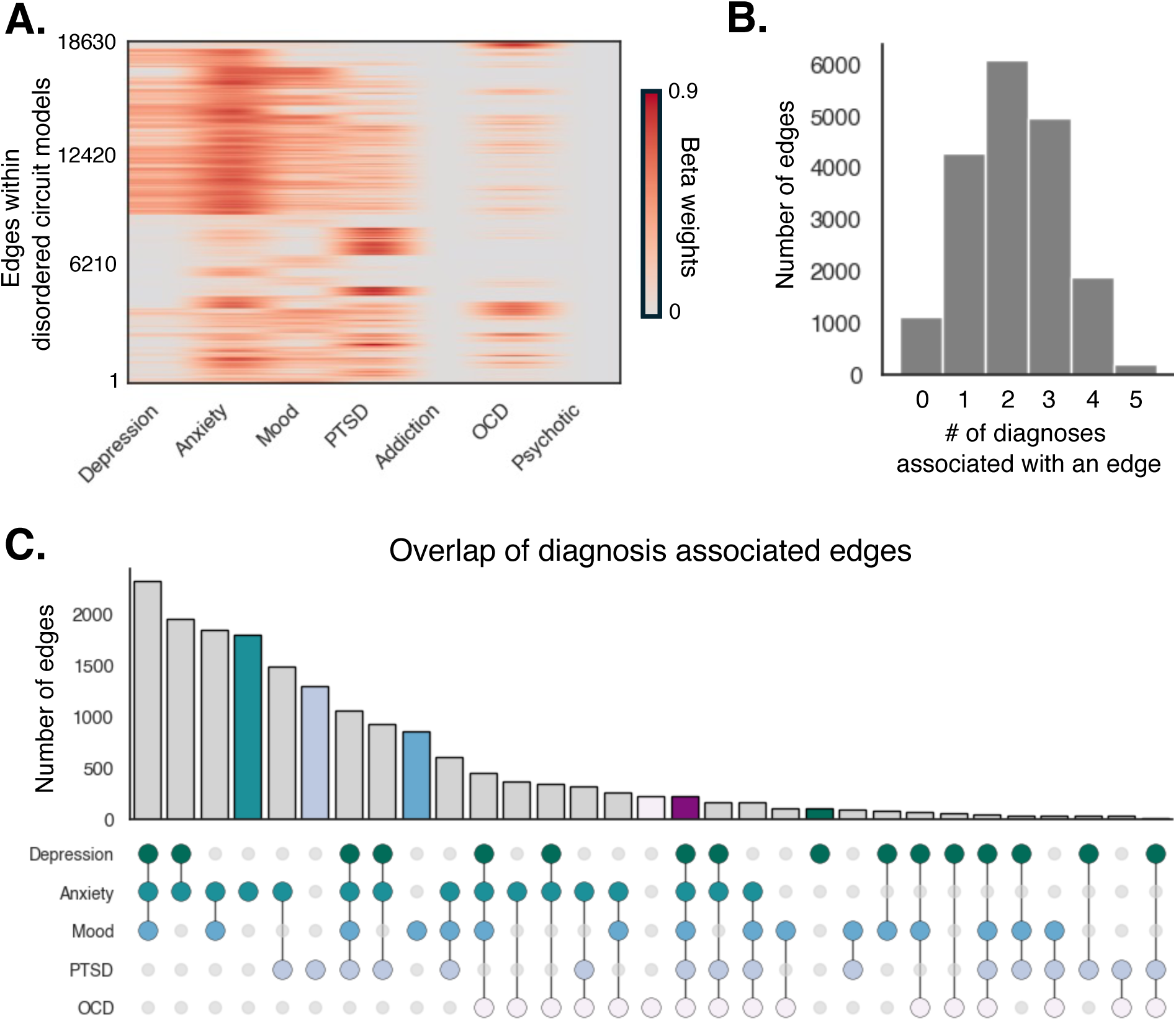
Relationships between disordered circuits and diagnostic categories. A) Multiple regressions on each edge in disordered circuit models yielded beta weights describing how strongly each edge was associated with each diagnosis. Edges that were not significantly associated with a diagnosis after correcting for multiple comparisons were assigned a beta weight of 0. B) Histogram showing the number of edges in the disordered circuit models associated with the number of diagnoses. C) A plot showing how many edges were associated with diagnostic comorbidity patterns and diagnostic specificity.

To determine whether disordered circuit models contained more diagnostic specific information than symptoms, we next examined whether individual symptoms were uniquely related to specific diagnostic categories. The multiple regression analyses described above were repeated to assess the relationship between each symptom and each diagnosis. Of the 24 symptoms assessed, none were specific to a single diagnostic category (**Supplemental Fig 5**). This highlights the extensive symptom comorbidity across diagnoses and suggests that disordered circuit models contain more diagnosis-specific information than symptoms alone. Furthermore, this suggests that individuals may arrive at symptoms shared between diagnoses via different neurobiological routes.

Although transdiagnostic symptoms did not show specificity to individual diagnostic categories, it was possible that disordered circuit models did because symptom networks encompassed a broader space and offered more degrees of freedom, instead of heterogeneity in edge dysfunction being the driving factor. Therefore, we examined whether disordered circuit models constructed solely on the complex shared/unique network space (and not weighted by edge-level dysfunction) also identified edges related to specific diagnoses. To test this, we repeated this analysis on disordered circuit models weighted only by symptom severity rather than by both symptom severity and edge dysfunction. This approach identified few diagnosis-specific edges, with only 431 edges unique to anxiety disorders and none that were unique to the other diagnostic categories. Instead, most edges were significantly related to all five diagnostic categories (5299/18630 or 28.44%; **Supplemental Fig 6**). Together, these results underscore the importance of incorporating information about individual-specific edge-level heterogeneity for understanding mechanistic differences in how similar symptoms arise in separate diagnostic categories.

Finally, localizations of the disordered circuits specific to each diagnostic category and shared across all 5 diagnoses were described in terms of the Yeo17 canonical networks^52^ (plus brainstem, cerebellum, and subcortex) (**Fig 5**). The top 3 canonical network pairs with the highest concentrations of disordered circuits specific to depression were FPCN A to DMN A, FPCN A to DAN A, and DMN A to DMN B. The top networks with circuits specific to anxiety diagnoses were within Limbic B, between Limbic B to brainstem, and DAN B to brainstem. The top networks with circuits specific to mood disorder diagnoses were between VAN A to FPCN A, Limbic B to FPCN A, and DAN A to DAN B. The top networks with circuits specific to PTSD were within DMN B, between DMN A to DMN B, and within brainstem. The top networks with circuits specific to OCD were between SM B to FPCN A, SM B to FPCN B, and within SM A. The top networks with circuits shared across all 5 diagnoses were between VAN B to DMN B, FPCN A to subcortex, and FPCN B to FPCN C. The top 5 canonical network pairs with the highest concentrations of disordered circuits specific to each diagnostic category and shared between all diagnoses are highlighted in **Supplemental Table 2**.

**Figure 5.**
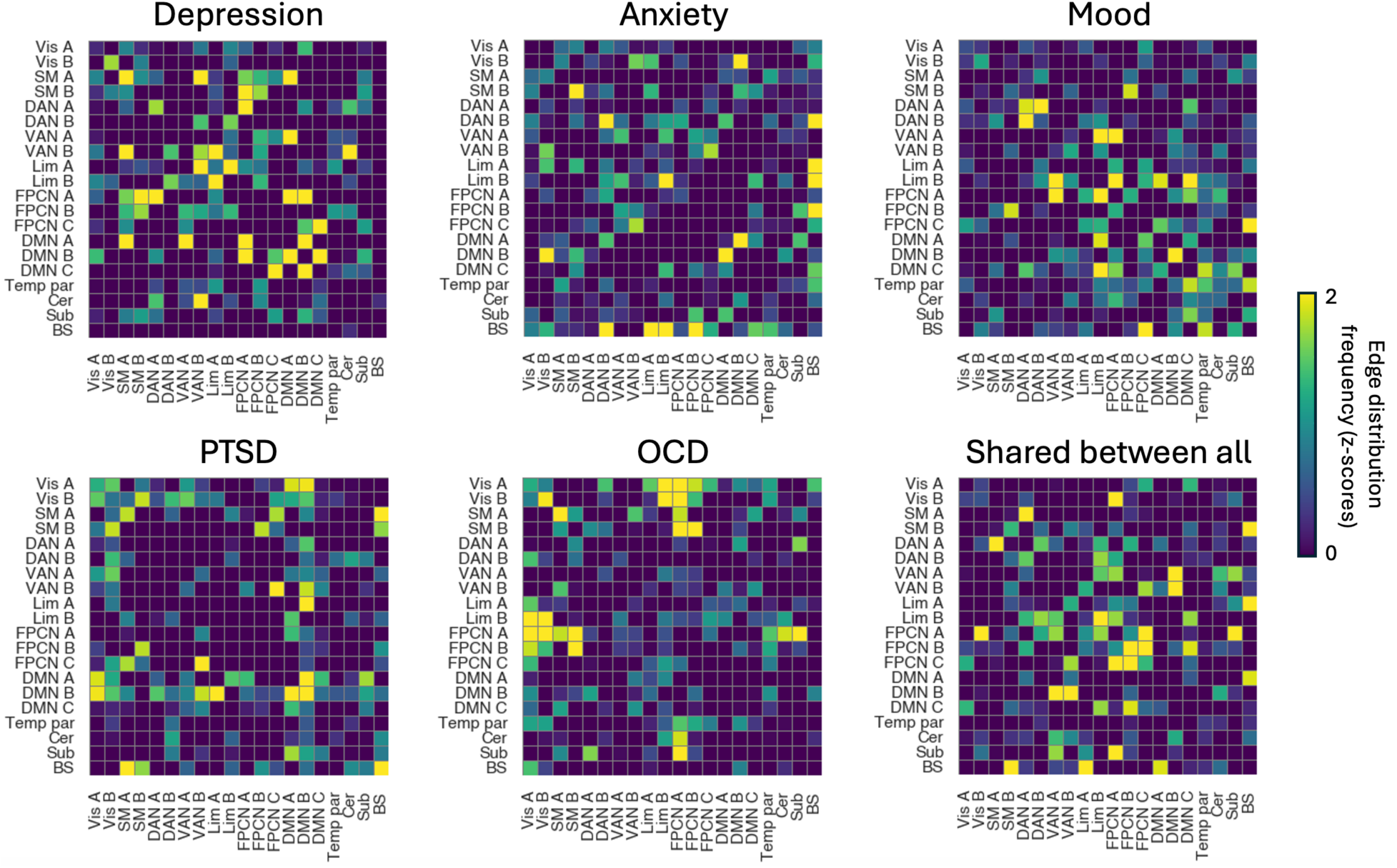
Network localization of edges specific to each diagnosis and shared between all diagnoses. Localizations of edges were highlighted by summing the number of times an edge specific to a diagnosis was distributed between each possible network-network pair and normalizing that number by the total number of network-network connections.

## Discussion

Identifying functional connectivity networks underlying transdiagnostic symptomatology holds promise to inform neurobiologically-guided precision psychiatry frameworks^17,53^. While the field has made substantial strides toward understanding the relevance of functional connectivity networks to psychopathology, the goal of yielding clinically-actionable insights from this research has yet to be realized^8,24,25^. In addition to the complexities of symptom heterogeneity within disorders^2,9,10^, this lack of clinical translation may also be driven by symptoms arising from heterogenous neural mechanisms^2,27,28^. As we show in this study, brain-wide functional connectivity networks linked to symptoms are not uniformly affected in a given individual. To account for this, we developed disordered circuit modeling, which characterizes individual-specific neural dysfunction patterns across multiple symptom networks. The disordered circuit models simultaneously accounted for a patient’s multivariate symptom profile, connectivity within the complex space of shared and unique symptom networks, and edge-level network heterogeneity. They contained information about overall diagnostic burden, co-morbid symptomatology patterns spanning multiple diagnostic categories, and information selectively associated with individual psychiatric diagnoses. Thus, identifying individualized disordered circuit modeling could provide a means through which connectivity-based models provide clinical utility within precision psychiatry settings.

### Leveraging network overlap for understanding disordered circuit locations

Previous work has demonstrated that network correlates of transdiagnostic symptoms exist in a complex web of overlapping and non-overlapping features^36,40–45^. According to a cornerstone hypothesis of the RDoC framework^15,16^, the locations of an individual’s disordered brain circuits should relate to their multivariate symptom profiles. As such, in this study, we sought to leverage the shared and unique symptom network space to gain a better understanding of the neurobiological basis of symptom comorbidity. We hypothesized that disordered circuits related to comorbid symptomatology should be distributed in areas shared between multiple symptom networks. Conversely, disordered circuits specific to a particular symptom should be distributed networks uniquely associated with that symptom. Tools for interrogating this have been underdeveloped to date. Our approach of aggregating multiple symptom networks into a single model represents a major step towards improving our understanding of the neurobiological basis of different combinations of symptom comorbidities. This approach can be extended to many different conditions in which symptom comorbidities are common to identify the shared neurobiological correlates. Such an endeavor could reveal novel network targets for psychiatric comorbidities.

### Capitalizing on heterogeneity in connectivity models

In demonstrating that substantial edge-level heterogeneity exists within network models of psychiatric symptoms, we provide evidence in support of the hypothesis that there are numerous neural routes to similar symptom presentations^2,27,28^. Furthermore, we show that the structure of this heterogeneity tracks with different diagnostic profiles. This suggests that differences in the neurobiology related to symptoms commonly shared across diagnostic categories, such as trouble falling asleep in major depression, generalized anxiety disorder, and PTSD^13,14^, may reflect variations in how that symptom arose in the first place. This phenomenon may shed light on why transdiagnostic network-based biomarkers have yet to be identified that are robust enough to reliably guide diagnostic and treatment strategies^8,24,25^. Therefore, studies seeking to identify network correlates of psychiatric symptoms with the intention of guiding precision medicine frameworks should be accounting for these heterogenous mechanisms that give rise to a symptom.

This finding also has major implications for guiding improvements to predictive network model generalizability. Building models that generalize to external data has been a top priority for advancing the field to the point where it can guide clinical decision making^54,55^. While efforts have been largely focused on determining optimal sample sizes and scanning time needed to create replicable network models of psychopathological measures^39,56^, heterogenous mechanisms underlying a symptom has not been effectively accounted for. Increasing the sample size in training data likely improves model generalizability because more subjects lead to a greater representation of the diverse neurobiological pathways linked to a symptom. Thus, adequately accounting for mechanistic differences in how symptoms arise in an individual may improve predictive accuracy in unseen data.

### Disordered circuits linked to specific diagnoses and comorbidity

It is well known that symptoms are highly co-morbid across diagnostic categories^11,12^. Reflecting this phenomenon, we showed that none of the symptoms we assessed were specifically linked to any one diagnostic category. However, many features within the disordered circuits were, suggesting that transdiagnostic symptomatology can stem from different neurobiological processes depending on a patient’s diagnostic profile. Importantly, these diagnostic specific network features identified from our data-driven approach align with evidence from other literature, including theoretical models and hypothesis-driven case-controlled studies. For example, a highly cited meta-analysis noted that major depression disorder was characterized by abnormal connectivity between the frontoparietal and dorsal attention networks, between the frontoparietal and default mode networks, and within the default mode network, which were the top 3 networks to which the disordered circuits specific to depression were localized^57^. Interestingly, this same meta-analysis also reported that a core feature of depression was abnormal connectivity within the frontoparietal network, but our results indicate that disordered circuits within the frontoparietal network tended to be linked to comorbid pathology and were not specific to depression. This highlights the utility of disordered circuit modeling for disentangling which circuits are specific to diagnostic categories and which are linked to comorbidity. Indeed, circuits within the frontoparietal network have been linked to psychiatric comorbidity elsewhere^42^. Additionally, our models showed that brainstem to limbic and within limbic network connectivity was specifically linked to anxiety disorders. These findings are consistent with extensive work in humans and other species highlighting that elevated communication between brainstem nuclei and limbic structures are core components of anxiety circuitry^58,59^. Furthermore, our findings specifically linking frontoparietal to limbic network connectivity to mood disorders like bipolar^60,61^, within default mode network connectivity to PTSD^62,63^, and somatomotor network dysfunction to OCD^64^ are also all consistent with the broader literature. Taken together, this suggests that disordered circuit modeling can effectively capture the plethora of neurobiological routes to transdiagnostic symptomatology at the individual level.

### Potential utilities

We envision that disordered circuit modeling will be useful across several lines of research relevant to the development of precision medicine frameworks. Circuit models have been leveraged to identify transdiagnostic biotypes that have been shown to effectively stratify patients into biologically-grounded subgroups that differentially respond to various forms of treatment^3,5,65^. These circuit models were defined *a priori* and thus did not fully encapsulate an individual’s total possible disordered circuitry. Using data driven approaches, like the one proposed in this study, could achieve this, and thus could provide more granular information to guide biotyping procedures. Better optimized biotyping pipelines may reveal more precise neuromodulation targets and improve treatment response predictions^54^. The inverse of this approach – that is, identifying well-preserved edges instead of disordered ones in an individual – could potentially reveal protective or compensatory neural mechanisms, such as those theorized to protect against cognitive decline coinciding with Alzheimer’s Disease pathology^66^. While there is behavioral evidence for neural compensation^67^, identifying circuit markers has remained elusive^68^. Much like how disordered circuit modeling capitalized on finding the most disordered circuits, this same pipeline could also identify the least disordered circuits and examine how these interface with the circuits most affected by pathology and networks supporting cognition. Finally, identifying links between disordered circuit patterns and multivariate genetic interactions revealed from psychiatric GWAS research^69^ would yield insights that bridge fields and provide a better understanding of the neurobiological underpinnings of circuit dysfunction.

### Limitations

This study had a few noteworthy limitations. First, while the predictive power we observed is on par with what is commonly seen in studies modeling self-reported symptoms (*ρ* = 0.15-0.2)^40^, the power is still low and leaves room for localization error in the initial network mapping steps. While the sample size here is large and transdiagnostic, generalizing the models to external datasets would improve confidence in the networks derived, but openly available datasets that provide item-level BSI data that are also harmonized with scanning protocols and population characteristics of the dataset we used do not exist. However, the purpose of this study was to demonstrate that variation within predictive symptom networks is meaningful and to capitalize on this variation, rather than to show that predictive networks are generalizable. Second, since the diagnostic category definitions and subjective symptom assessments are subject to response bias^70^, benchmarking these models against a ‘ground truth’ is unfeasible. Finally, our inability to identify circuits related to some diagnoses, such as psychoticism, was likely because there was only one modellable item related to psychotic symptoms and patients with psychotic diagnoses were relatively underrepresented in this dataset.

## Methods

### Participants

The sample reported in this study includes data from 317 demographically diverse participants (age range = 18-73 years; mean age = 31.17 +/- 11.17 years, 174 females and 143 males) with >50 minutes of high-quality functional MRI data. This dataset is described in greater detail by Samardzija et al. (2025)^20^ and is openly available on OpenNeuro. Participants were recruited from advertisements broadly distributed throughout the New Haven community and referrals from Yale clinics. This transdiagnostic population experienced a wide range of symptoms and symptom severities, often with multiple psychiatric diagnoses (**Supplemental Table 1**). All participants provided written informed consent in accordance with a protocol approved by the Yale IRB.

### Imaging acquisition protocol

The imaging data was collected at Yale on a 3T Siemens Prisma scanner with a 64- channel head coil. A high-resolution T1-weighted MPRAGE (TR= 2,400 ms, TE = 1.22 ms, flip angle = 8°, voxel size 1 mm^3^) was acquired for each participant. The functional data were obtained using a multiband EPI sequence (TR = 1,000 ms, TE = 30 ms, flip angle = 55°, slice thickness = 2 mm, multiband factor = 5). A total of 6 task and 2 resting-state runs were acquired, each lasting 6 minutes and 49 seconds. The first and last functional scans were resting-state runs. The six task runs were comprised of a working memory (n-back^71^), an inhibition (stop signal task^72^), a decision making (card guessing^73^), an emotional perception (reading the mind in the eyes^74^), a continuous performance attention task (the gradCPT^75^), and a movie watching condition^20^. The task order was randomized and counterbalanced across participants. We ensured that participants understood task instructions and received practice. Analyses were restricted to participants who completed all fMRI scan runs.

### Functional MRI data processing

Skull stripping of the structural scans was done using an optimized version of the FMRIB’s Software Library (FSL) pipeline^76,77^. Motion correction was done using SPM12^78^. Nonlinear registration of the MPRAGE to the MNI template was performed using BioImage Suite^79^, and linear registration of the functional to the structural images was done through a combination of FSL^80^ and BioImage Suite. The remaining preprocessing steps were performed in BioImage Suite, including global signal regression, high-pass and low-pass filtering, and regression of motion parameters. All registered data were visually examined to ensure whole-brain coverage, adequate registration, and the absence of artifacts or other quality issues. Participants were included in the study if they completed all eight-fMRI scan runs and had a grand mean frame-to-frame displacement of less than 0.15 mm and a maximum mean frame-to-frame displacement of less than 0.2 mm. Two participants were scanned with a slightly shorter scanning time (25 seconds shorter) and were included in the dataset.

The Shen 268 node atlas^81^ was applied to the preprocessed data, parcellating it into 268 functionally coherent nodes. Pearson correlations of the time series between all node pairs were computed and subsequently z-transformed to generate 8 functional connectivity matrices for each participant. The functional connectivity matrices were averaged across all runs for each participant. Aggregating data from different scanning conditions accounted for patterns of task-evoked brain activity that may differentially enhance symptom predictions^82^. Analyses and visualizations were conducted using python.

### Psychiatric symptom assessments

A variety of psychiatric symptoms were assessed using the Brief Symptom Inventory (BSI)^50^ immediately following the MRI scans (same day). This self-reported symptom assessment was composed of 50 items that probed how often within the past two weeks the participant experienced symptoms related to depression, anxiety, phobic anxiety, psychoticism, paranoid ideation, interpersonal sensitivity, hostility, somatization, obsession-compulsion, guilt, trouble sleeping, and poor appetite. Importantly, the BSI contains items related to different aspects of similar symptoms (e.g., multiple items related to depression symptoms), enabling us to better capture heterogenous symptom profiles. A few participants did not complete all clinical assessment scales. One participant did not complete any of the BSI assessments and was therefore excluded from disordered circuit modeling. Two participants did not provide a response to one item, and one participant did not provide responses to two items. Because these three individuals provided data for >90% of modellable items, they were included in disordered circuit modeling.

### Categorical diagnostic information

Participants were also assessed for whether they met criteria for a variety of major psychiatric disorders in the DSM-5 and ICD-10 using the Mini-International Neuropsychiatric Interview (MINI)^51^. Assessments were conducted on the same day as the fMRI scanning. Participants who met criteria for current, recurrent, or past major depressive episode were noted as meeting criteria for a depression diagnosis. Participants were marked as meeting criteria for an anxiety disorder if they met criteria for any of the anxiety disorders indexed by the MINI (generalized anxiety disorder, social anxiety, phobias, or a panic disorder). Obsessive-compulsive disorder (OCD) was considered separately because it is not classified as an anxiety disorder in the DSM-5. Participants were noted as having a mood disorder if they met criteria for either bipolar, hypomanic/manic episodes, or an unspecified mood disorder. Participants were noted as having post-traumatic stress disorder (PTSD) if they met criteria for either past or present PTSD. Participants were considered to have an addiction disorder if they met criteria for either alcohol or substance use disorders. Finally, participants were noted as having a psychotic disorder if they met criteria for either a psychotic disorder or bipolar or mania with psychotic features.

### Identifying symptom networks

Functional connectivity networks related to symptoms were identified using CPM. This method has been extensively described previously^29,30^. Briefly, we utilized a 10-fold cross validated predictive model to identify the predictive utility of edges that were significantly correlated (*p* < 0.05) with behavior in the training folds after controlling for age and sex using partial correlations. Prediction strength was determined by comparing the observed symptoms to the predicted symptoms using Spearman correlations. Modeling was repeated 1000 times to obtain a distribution of predictions. Significance was determined by comparing the observed prediction strengths to a null distribution generated from running CPM on 1000 randomly permuted observations. *P*-values were obtained by calculating the proportion of null predictions that exceeded the median value from the actual prediction strength distribution. All *p*-values were FDR corrected for multiple comparisons using the Benjamini-Hochberg procedure. Network correlates for each measure were derived by identifying the edges that were predictive in >50% of permutations. The resulting networks were binarized, with a ‘1’ or ‘0’ indicating an edge’s network membership. Symptom network similarity was assessed using the Dice similarity metric.

### Assessing heterogeneity within symptom networks

To examine how the edges within CPM-derived symptom network models were heterogeneously disordered between individuals, linear models describing the relationship between connectivity and symptomatology for each edge within the symptom networks were calculated. From these edge-specific brain-behavior models, residuals were obtained for each edge and participant. Edge residuals were converted to z-scores relative to the fit-line, and the signs were flipped for edges with a negative brain-behavior model slope to facilitate comparisons between edges with positive brain-behavior model slopes. By residualizing edge connectivity, we were able to similarly scale edge dysfunction across a range of symptoms. Since the CPM models suggest that higher connectivity is related to worse symptomatology, then an individual’s edges with higher residual scaled values should be contributing most to their symptoms. Interindividual similarities and differences in how scaled edge connectivity was distributed within each symptom network was assessed using Person’s correlations between all possible pairs of participants for every symptom network. Permutation tests were utilized to determine whether scaled edge connectivity was randomly distributed throughout symptom networks, or if there was systematic structure in how they were distributed between individuals. Specifically, the kurtosis of the distribution of correlation coefficients describing interindividual edge similarities was computed for each symptom network. Then, 1000 randomly generated null distributions were obtained by shuffling each individual’s scaled edge connectivity values within the symptom network and recomputing pairwise correlations between all individuals. A higher observed kurtosis value than random would indicate that individuals tended to display greater correlations and anticorrelations in their edge residuals’ locations than expected by chance. So, the observed interindividual edge similarity distribution kurtosis was compared to the null distributions to obtain *p*-values. Specifically, the number of null kurtosis values that exceeded the median observed kurtosis value divided by the number of iterations of random shuffling (1000) yielded the *p*-values. This was repeated separately for each symptom network and *p*-values were FDR corrected for multiple comparisons.

### Constructing disordered circuit models

Disordered circuit models were designed to incorporate information from an individual’s multivariate symptom profile, the architecture of the shared and unique symptom network space, and the heterogeneity within symptom networks. To achieve this, node x node binarized masks for each symptom network were vectorized, with the diagonal and lower triangle being removed (due to the networks’ symmetry). The binarized network mask vectors from all symptom networks were then stacked in a symptom x edge matrix. Then, for every individual, each edge within CPM-derived binarized symptom network masks were separately weighted by their residual scaled edges to encompass edge-level heterogeneity. Prior to weighting, the residual scaled edges, which were initially z-scored, were then scaled from 1-10 to avoid multiplying by negative numbers. Then, the edge weighted symptom networks were weighted by the scaled symptom score for each individual to highlight each participant’s edges that contribute most to the CPM models at the group level. Similarly, symptoms were first z-scored and then scaled from 1-10 before weighting. Once each symptom network was doubly weighted by edge-level dysfunction and symptom scores, the weighted edges were summed across all symptom networks (the rows). By doing this, the edges that were consistently associated with poor symptomatology were emphasized. After repeating this procedure for each participant, each edge was z-scored across all participants to facilitate comparisons between edges that appeared in multiple symptom networks and edges that appeared in one (or few). Edges that were not predictive of any symptoms (i.e., did not appear in any symptom networks) were not used for further analyses. In total, disordered circuit values were derived for 18,630 out of the 35,778 total edges in the functional connectome. All edge z-scores were offset by the minimum edge z-score to ensure that all values were positive to facilitate interpretations of the data. The resulting vectors represented the disordered circuits for each participant. Stability was assessed by performing 1000 iterations of a split-half resampling procedure. For each iteration, participants were randomly divided into two equal halves. All parameters required to construct disordered circuit vectors – including symptom normalization, edgewise regression models, residual standardization, and final edgewise normalization – were estimated using one half of the sample and then applied exclusively to the held-out half. The procedure was then repeated with the training and test halves reversed, such that every participant received an out-of-sample disordered circuit vector generated without contributing to the estimation of the underlying model parameters. Stability was quantified as the Pearson correlation between each participant’s out-of-sample disordered circuit vector and the corresponding circuit vector constructed on the whole sample.

### Linking disordered circuit features to diagnostic categories

To identify the disordered circuits that were linked to individual diagnostic categories and to co-morbid diagnoses, multiple linear regression models for each edge were utilized where the diagnostic categories were set as the dependent variables and the edge’s disordered circuit magnitude was set as the independent variable. All derived *p*-values were FDR corrected for multiple comparisons. The Yeo17 canonical networks^52^ (plus brainstem, cerebellum, and subcortex) were used to describe where the disordered circuits specific to each diagnostic category and shared across all 5 diagnoses were localized to. To do so, binarized adjacency matrices representing the disordered circuits specific to each diagnosis and shared between all diagnoses were constructed. The number of times that disordered circuit edges were connected to each canonical network pair was summed. These sums were then divided by the total number of edges distributed between canonical networks to adjust for network size. This yielded 20x20 matrices representing how disordered circuits were distributed across networks. These matrices were z-scored to enable the visualizations to be scaled the same across diagnostic categories.

## Acknowledgements

We would like to thank Hilary Blumberg and Ilan Harpaz-Rotem for helpful conversations.

## Funding statement

This work was supported by funding from the National Institutes of Health (MH121095, MH138347, and EB034720 to R.T.C.). A.J.S. is also supported by the National Science Foundation’s Graduate Research Fellowship Program.

## Data availability

The demographic, behavioral, and fMRI utilized in this study is publicly available^20,83^ (<u>10.18112/openneuro.ds007286.v1.0.5</u>). The prediction strength values and derived network masks for all BSI items have been archived on Zenodo and can be found at: https://doi.org/10.5281/zenodo.23577041.

## Code availability

The code used in all the main analyses conducted in this study has been archived on Zenodo and can be found at: https://doi.org/10.5281/zenodo.23577041. All dependency files required to run this code and relevant input data are also provided in this repository.

## Competing interests

G.S. has served as a consultant or scientific advisory board member to Axsome Therapeutics, Biogen, Biohaven Pharmaceuticals, Boehringer Ingelheim International, Bristol-Myers Squibb, Clexio, Cowen, Denovo Biopharma, ECR1, EMA Wellness, Engrail Therapeutics, Gilgamesh, Janssen, Levo, Lundbeck, Merck, Navitor Pharmaceuticals, Neurocrine, Novartis, Noven Pharmaceuticals, Perception Neuroscience, Praxis Therapeutics, Sage Pharmaceuticals, Seelos Pharmaceuticals, Vistagen Therapeutics and XW Labs; and received research contracts from Johnson & Johnson (Janssen), Merck and Usona. G.S. holds equity in Biohaven Pharmaceuticals and is a co-inventor on a US patent (8,778,979) held by Yale University and a co-inventor on US provisional patent application no. 047162-7177P1 (00754), filed on 20 August 2018 by Yale University Office of Cooperative Research. Yale University has a financial relationship with Janssen Pharmaceuticals and may receive financial benefits from this relationship. The University has put multiple measures in place to mitigate this institutional conflict of interest. Questions about the details of these measures should be directed to Yale University’s Conflict of Interest office. V.H.S. has served as a scientific advisory board member to Takeda and Janssen. The remaining authors declare no competing interests.

## Author contributions

Based on <u>CRediT roles</u> Conceptualization; A.J.S., R.T.C. Data curation: A.J.S., S.I., Anja S., F.P., K.Y.T. Formal analysis; A.J.S. Funding acquisition; R.T.C Investigation: A.J.S. Methodology; A.J.S., R.T.C. Project administration; F.T., M.Q., J.A. Resources; R.K., G.S., S.W.W., V.H.S. Software; A.J.S., X.S. Supervision; R.T.C. Visualization; A.J.S. Roles/Writing - original draft; A.J.S., R.T.C Writing - review & editing. All authors

## Notes

https://doi.org/10.5281/zenodo.23577041

### Summary of Updates

Figure 2 was revised, along with the results and methods that correspond to the analyses presented in Figure 2. The abstract, introduction, and discussion were revised to place greater emphasis on the clinical implications of this study (how there are multiple neurobiological routes to the same symptoms) and to frame the methodological approach we developed to modeling personalized disordered circuits as a means to understand this clinically relevant problem.

