## Supplemental materials for "Personalized disordered circuit modeling reveals diverse neurobiological routes to transdiagnostic symptoms"

**Supplemental Table 1**. Number of participants in each diagnostic category in the MINI grouped by diagnostic label used in the main text.

| Diagnostic label assigned in main text | MINI diagnosis and n |
|---|---|
| Depression | Major depressive episode = 107<br>Major depressive episode (past) = 105<br>Major depressive episode (current) = 35<br>Major depressive episode (recurrent) = 54 |
| Anxiety | Generalized anxiety disorder = 44<br>Panic disorder = 33<br>Agoraphobia = 14<br>Social anxiety disorder = 13<br>Agoraphobia = 14 |
| Mood | Hypomanic = 6<br>Hypomanic (past) = 5<br>Bipolar disorder = 25<br>Bipolar + psychotic = 5<br>Bipolar (past) = 4<br>Bipolar (current) = 2<br>Bipolar (single episode) = 2<br>Manic episode = 23<br>Manic episode (past) = 23<br>Manic episode (current) = 1<br>Manic episode + psychotic = 1<br>Mood (unspecified) = 7<br>Mood (unspecified, current) = 4<br>Mood (unspecified, lifetime) = 6<br>Mood (unspecified) + psychotic = 8<br>Panic = 33<br>Panic (lifetime) = 31<br>Panic (current) = 12 |
| PTSD | PTSD (total) = 22<br>PTSD (current) = 1<br>PTSD (past) = 20 |
| Addiction | Alcohol use disorder (total) = 27<br>Alcohol use disorder (mild) = 14<br>Alcohol use disorder (moderate) = 6<br>Alcohol use disorder (severe) = 5<br>Alcohol use disorder (remission) = 1<br>Substance use disorder (total) = 31<br>Substance use disorder (mild) = 16<br>Substance use disorder (moderate) = 6<br>Substance use disorder (severe) = 11<br>Substance use disorder (early remission) = 2 |
| OCD | Obsessive-compulsive disorder (OCD) = 14 |
| Psychotic | Psychotic = 8<br>Psychotic (current) = 6<br>Psychotic (lifetime) = 2<br>Bipolar + psychotic = 5<br>Manic episode + psychotic = 1<br>Mood (unspecified) + psychotic = 8 |
| Healthy controls | No diagnoses = 170 |

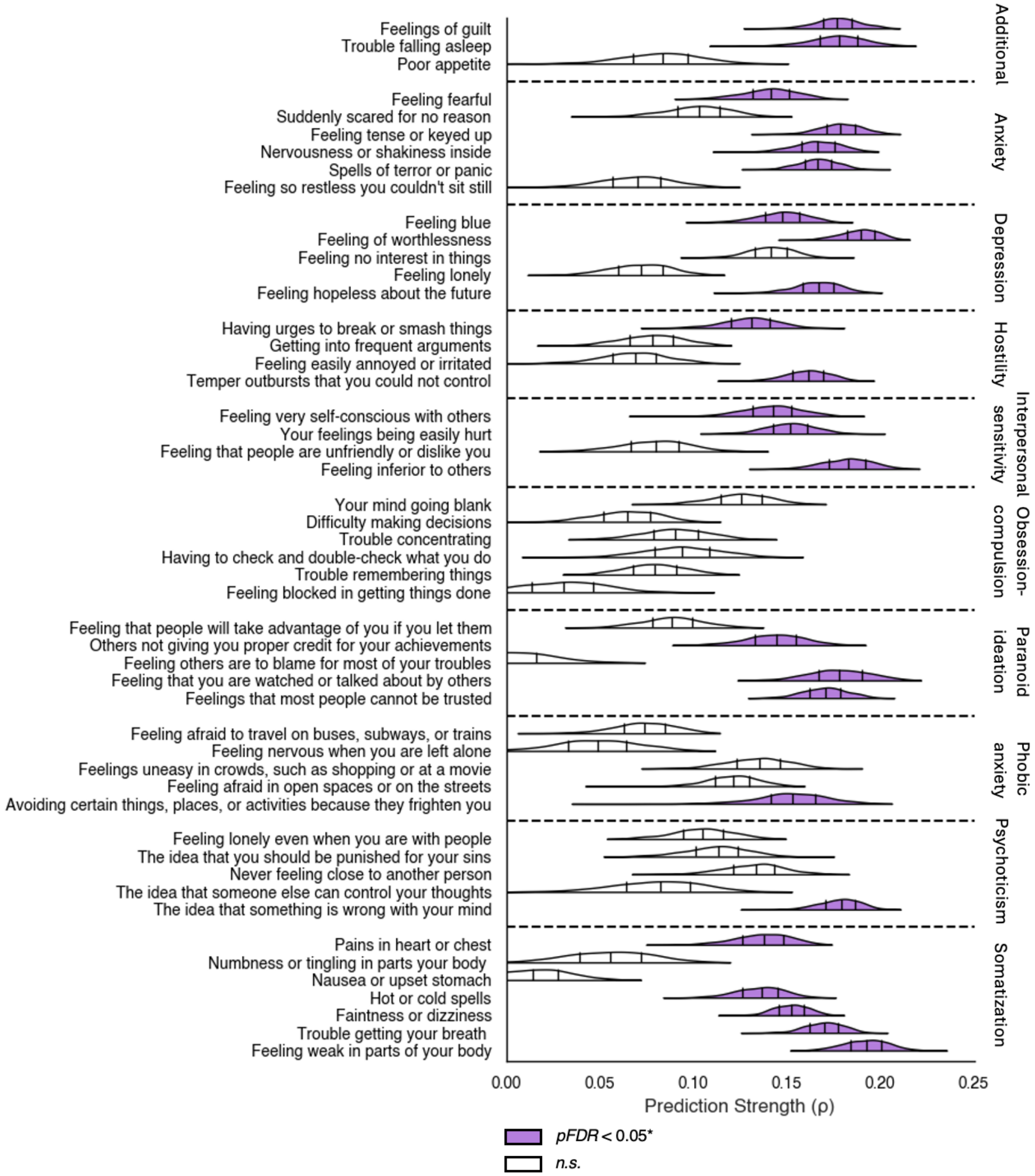


**Supplemental Figure 1**. Prediction strength of all items in the BSI.

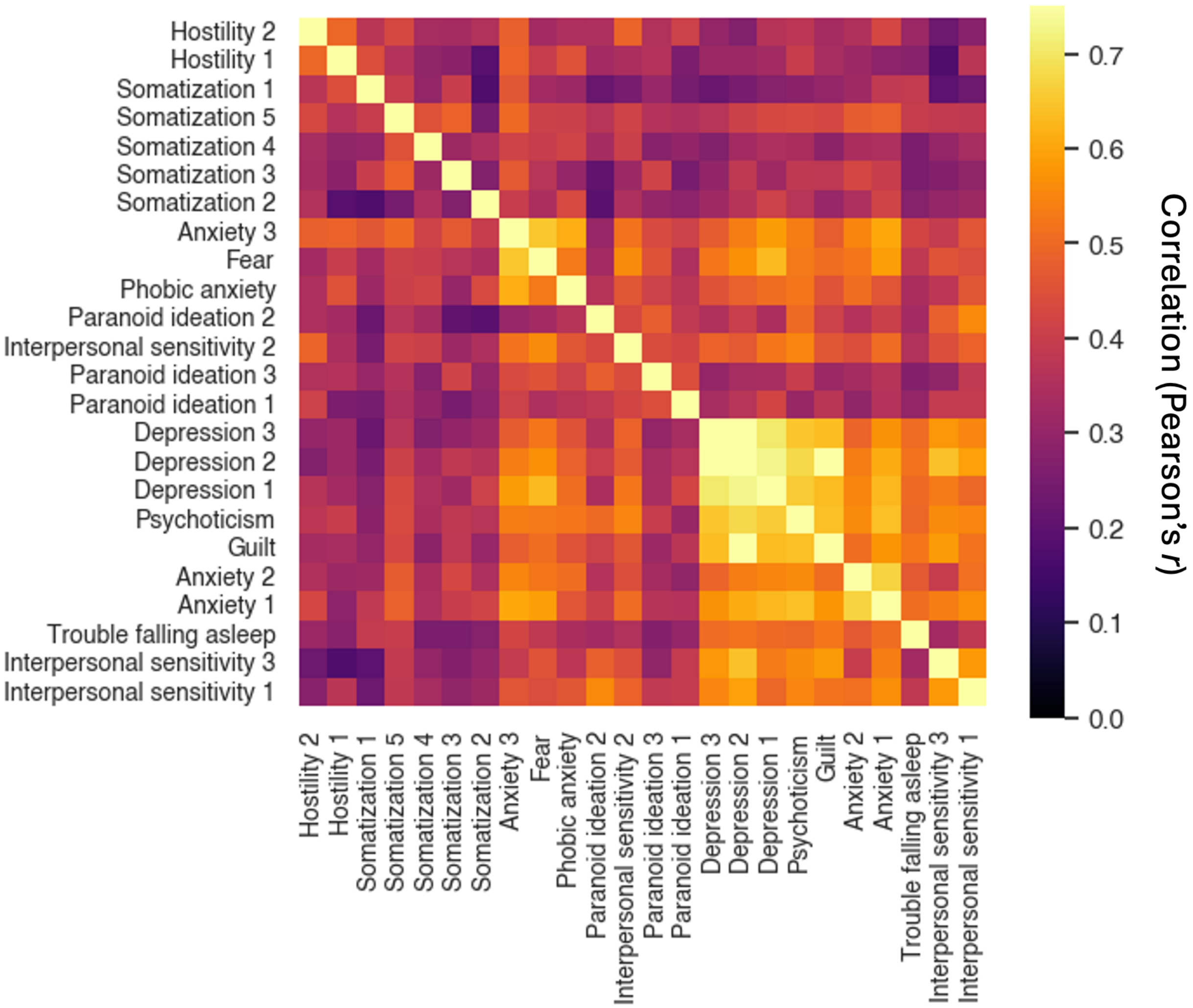


**Supplemental Figure 2.** Correlations between symptom scores. Symptom scores were sorted according to network similarity, as determined via agglomerative hierarchical clustering.

**Supplemental Table 2**. Summary statistics describing the interindividual similarities in scaled edge connectivity in each symptom network. Kurtosis values of the interindividual similarity distributions and *p*-values from comparing the observed kurtosis to null distributions' kurtosis values were derived from 1000 iterations of randomly shuffling each individual's scaled edge connectivity values and then computing null interindividual similarities. *P*-values were one-sided.

| Symptom network | Interindividual edge similarities (median *r* +/- standard deviation) | Interindividual edge similarity distribution kurtosis | *P*-values describing chance that the kurtosis was not random |
|---|---|---|---|
| Guilt | -0.005 +/- 0.064 | 0.395 | < 0.001 |
| Trouble falling asleep | -0.005 +/- 0.064 | 0.404 | < 0.001 |
| Fear | -0.005 +/- 0.063 | 0.491 | < 0.001 |
| Anxiety 1 | -0.005 +/- 0.065 | 0.381 | < 0.001 |
| Anxiety 2 | -0.004 +/- 0.070 | 0.385 | < 0.001 |
| Anxiety 3 | -0.005 +/- 0.060 | 0.572 | < 0.001 |
| Depression 1 | -0.004 +/- 0.063 | 0.455 | < 0.001 |
| Depression 2 | -0.005 +/- 0.062 | 0.477 | < 0.001 |
| Depression 3 | -0.005 +/- 0.064 | 0.478 | < 0.001 |
| Hostility 1 | -0.005 +/- 0.059 | 0.366 | < 0.001 |
| Hostility 2 | -0.005 +/- 0.057 | 0.410 | < 0.001 |
| Interpersonal sensitivity 1 | -0.005 +/- 0.070 | 0.339 | < 0.001 |
| Interpersonal sensitivity 2 | -0.005 +/- 0.065 | 0.466 | < 0.001 |
| Interpersonal sensitivity 3 | -0.005 +/- 0.064 | 0.356 | < 0.001 |
| Paranoid ideation 1 | -0.005 +/- 0.066 | 0.334 | < 0.001 |
| Paranoid ideation 2 | -0.005 +/- 0.070 | 0.344 | < 0.001 |
| Paranoid ideation 3 | -0.005 +/- 0.061 | 0.287 | < 0.001 |
| Phobic anxiety | -0.005 +/- 0.062 | 0.289 | < 0.001 |
| Psychoticism | -0.005 +/- 0.067 | 0.436 | < 0.001 |
| Somatization 1 | -0.005 +/- 0.059 | 0.378 | < 0.001 |
| Somatization 2 | -0.005 +/- 0.062 | 0.423 | < 0.001 |
| Somatization 3 | -0.005 +/- 0.059 | 0.415 | < 0.001 |
| Somatization 4 | -0.005 +/- 0.059 | 0.390 | < 0.001 |
| Somatization 5 | -0.005 +/- 0.062 | 0.309 | < 0.001 |

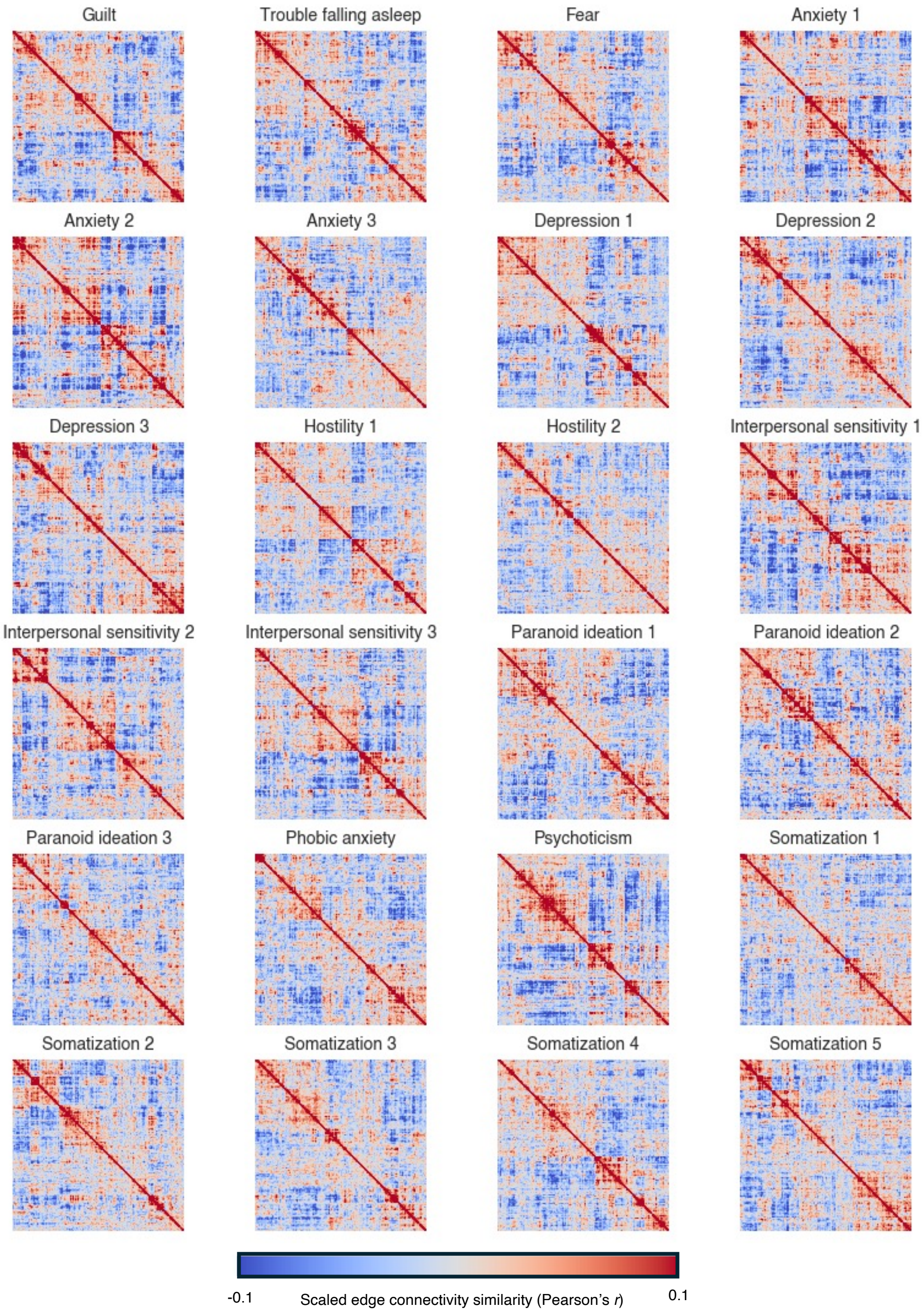


**Supplemental Figure 3.** Participant x participant correlation matrices showing how similar individuals were in how their edge connectivity was distributed throughout symptom networks. For visualization, participants were ordered based on similarity as determined by an agglomerative hierarchical clustering algorithm.

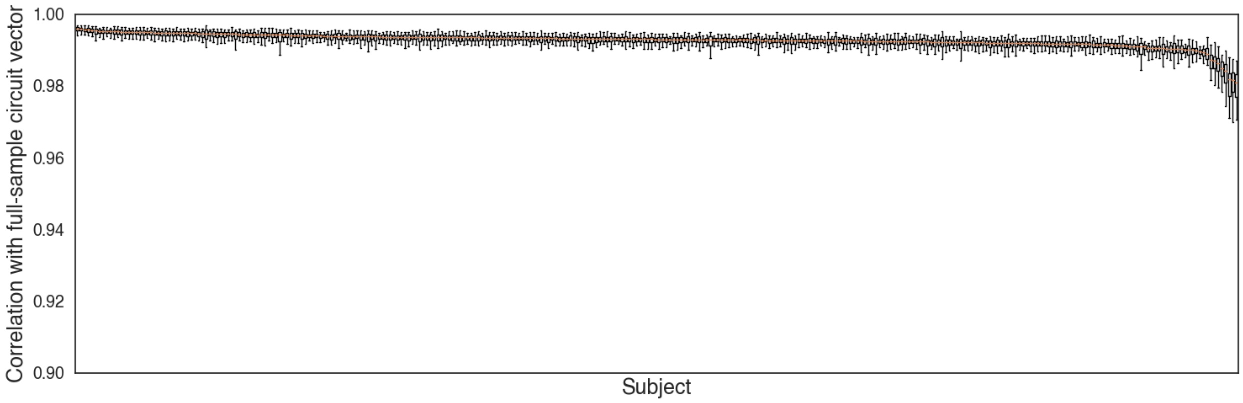


**Supplemental Figure 4.** Disordered circuit model stability. Each individual's disordered circuit vector median correlation to their resampled disordered circuit vector is displayed after 1000 iterations of split-half resampling. Subjects (x-axis) are sorted from left to right in order of descending median stability.

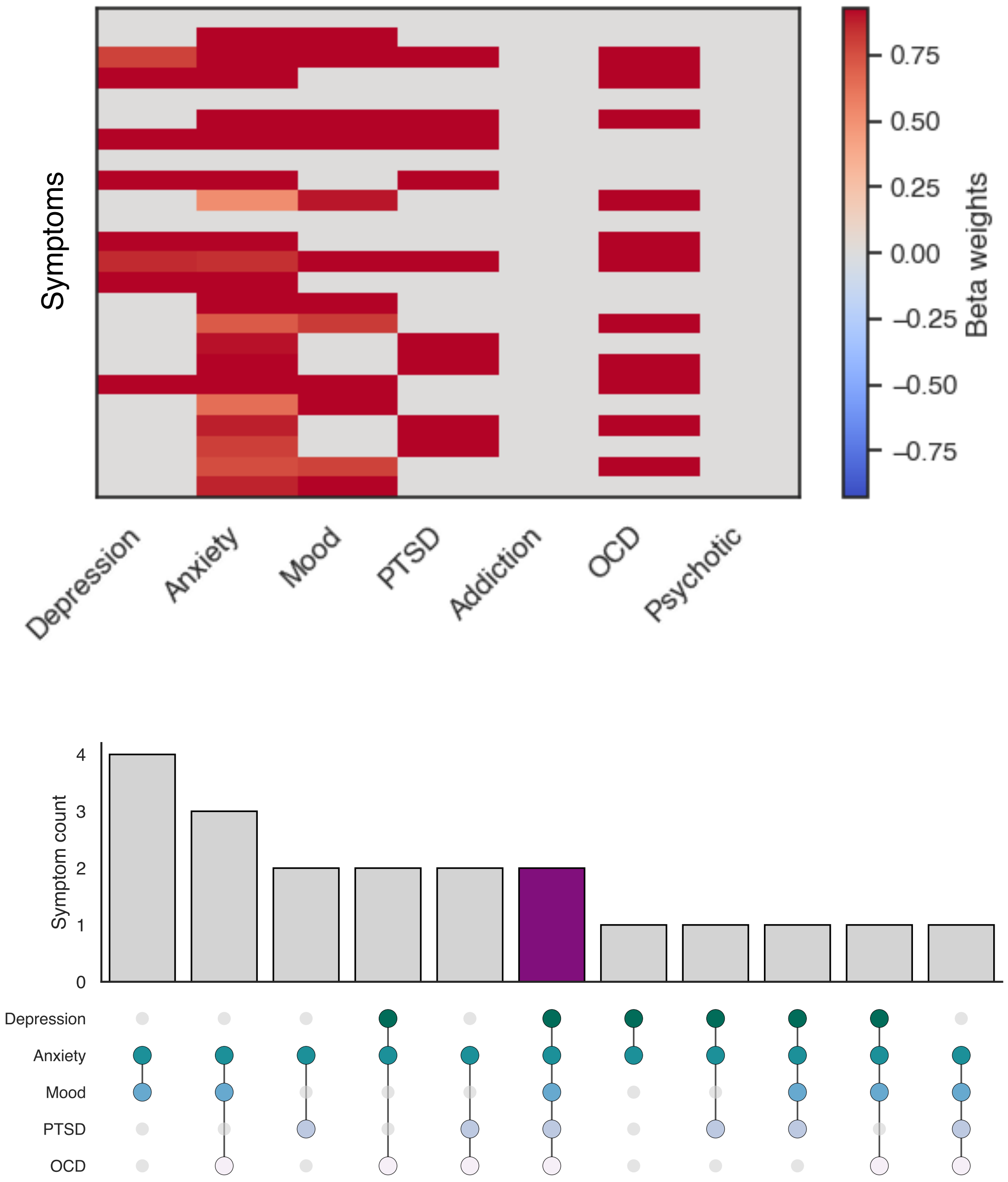


**Supplemental Figure 5. Shared symptoms across diagnostic categories.** Beta weights are displayed to highlight the strength of relationships between symptoms and diagnostic categories (top). Beta weights were only displayed for symptoms that were significantly associated ($p$ < 0.05) with diagnoses after correcting for multiple comparisons (*FDR*). No symptoms were uniquely associated with a single diagnosis (bottom).

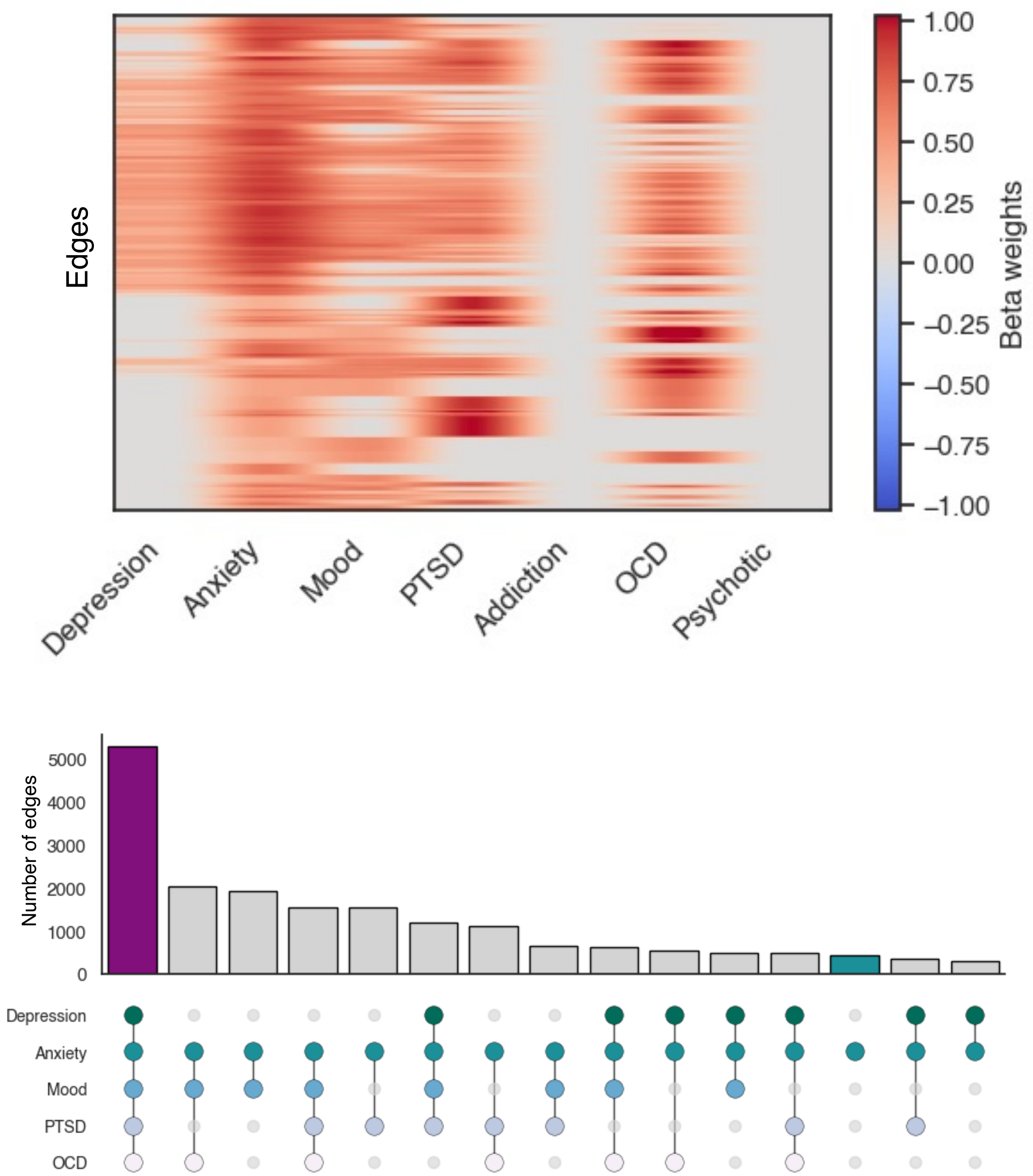


**Supplemental Figure 6. Mapping edges in disordered circuit models to diagnoses when edge heterogeneity within symptom networks was left unaccounted for.** Beta weights are displayed to highlight the strength of relationships between edges and diagnostic categories (top). Beta weights were only displayed for edges that were significantly associated ($p < 0.05$) with diagnoses after correcting for multiple comparisons (*FDR*). Edges were most commonly associated with 5 diagnoses when disordered circuit models were not weighted by edge heterogeneity.

**Supplemental Table 3.** Top 5 canonical network pairs with the highest concentrations of disordered circuit features specific to each diagnostic category and shared across all diagnoses.

| Depression | |
|---|---|
| FPCN A | DMN A |
| DAN A | FPCN A |
| DMN A | DMN B |
| SM A | VAN B |
| VAN B | Cer |
| Anxiety | |
| Lim B | Lim B |
| Lim B | BS |
| DAN B | BS |
| DAN B | DAN B |
| Lim A | BS |
| Mood | |
| VAN A | FPCN A |
| Lim B | FPCN A |
| DAN A | DAN B |
| DMN B | DMN B |
| VAN A | Lim B |
| PTSD | |
| DMN B | DMN B |
| DMN A | DMN B |
| BS | BS |
| SM A | BS |
| VAN B | FPCN C |
| OCD | |
| SM B | FPCN A |
| SM A | SM A |
| SM B | FPCN B |
| Vis B | Lim B |
| Vis B | FPCN A |
| Shared between all 5 diagnoses | |
| VAN B | DMN B |
| FPCN A | Sub |
| FPCN B | FPCN C |
| Lim A | BS |
| Vis B | FPCN A |
